# Integer topological defects organize stresses driving tissue morphogenesis

**DOI:** 10.1101/2020.06.02.129262

**Authors:** Pau Guillamat, Carles Blanch-Mercader, Karsten Kruse, Aurélien Roux

**Author notes:** These authors contributed equally to this work (Pau Guillamat, Carles Blanch-Mercader).

## Abstract

Tissues acquire their function and shape via differentiation and morphogenesis. Both processes are driven by coordinating cellular forces and shapes at the tissue scale, but general principles governing this interplay remain to be discovered. Here, we report that self-organization of myoblasts around integer topological defects, namely spirals and asters, triggers localized differentiation and, when differentiation is inhibited, drives the growth of cylindrical multicellular protrusions. Both localized differentiation and growth require specific stress patterns. By analyzing the experimental velocity and orientation profiles through active gel theory, we show that integer topological defects can concentrate compressive stresses, which we measure by using deformable pillars. Altogether, we envision topological defects as mechanical organizational centers that control differentiation and morphogenesis to establish tissue architecture.

## Main text

Morphogenesis establishes shapes of tissues during development. It comprises a wide variety of dynamic force-generating processes, founded on actin-based cell contractility and motility(*1*). Tissue folding, for example, can be driven by apical constriction(*2*). On the other hand, most cells in living tissues are polarized, featuring anisotropic distribution of their constituents. At the single cell level, polarity is usually coupled to directional motility and anisotropic contractility(*3*, *4*). At the tissue level, forces and polarities of many cells need to be coordinated to generate the stress patterns required for creating specific tissue architectures.

One usual marker of cell polarity is shape anisotropy(*1*). Akin to elongated molecules in liquid crystals(*5*), elongated cells can self-organize into patterns featuring long-range orientational order (*6*–*9*). Orientational fields often present topological defects, where the orientational order is ill-defined. Still, they imply very specific orientational configurations around their cores(*5*). In active systems – driven by internal energy-consuming processes – topological defects entail characteristic flow and stress patterns that depend on the defects’ topological strength s, which indicates the rotation experienced by the orientational field along a path encircling a defect’s core(*10*). In particular, the dynamics associated with half-integer defects (s=±1/2) – loops and triradii – have been thoroughly studied both experimentally(*10*–*12*) and theoretically(*13*, *14*). Importantly, in cell monolayers, the position of half-integer defects correlates with biologically-relevant processes such as cell extrusion(*15*) or changes in cell density(*16*).

Nevertheless, the role of integer defects (s=±1) – whorls, asters or vortices - in living systems is unknown. Yet, integer defects abound in nature(*17*), mostly because of their symmetry, as structural organizers of plants’ and animals’ body plans: spiraling organization of leaves or astral arrangement of spikes on the sea urchin shell are few examples. Thus, integer defects may also play essential roles during establishment of tissues’ architecture. Indeed, their position correlates with the establishment of the body plan of hydra during regeneration *in vivo*(*18*). *In vitro*, formation of cellular integer defects has been enforced by imposing the orientational field externally(*19*–*21*). Nevertheless, whether the active mechanics of integer defects is involved in modelling tissues remains unexplored.

To generate cellular integer topological defects we used C2C12 myoblasts, which not only are elongated cells but can differentiate and fuse to form myotubes, the precursors of skeletal muscle(*22*). Mechanical stimuli can induce differentiation(*23*, *24*). Consistently with previous studies(*8*, *25*), physical interaction between C2C12 cells resulted in collective alignment and, subsequently, in the emergence of long-range orientational order (Fig.1A, Fig.S1 and Movie S1). Also, as previously reported(*16*, *25*), orientational order decreased locally at places of frustrated cell arrangements, which constituted topological defects s=±½ (Fig.1A and Methods). After confluence, the spatial nematic correlation distance, *ξ*_*nn*_, reached a plateau at 190±10μm (mean±SE, as anywhere in this work, Fig.1B, Fig.S1), setting a characteristic length scale coinciding with inter-defect spacing (Fig.1B, inset, and Methods).

**Figure 1.**
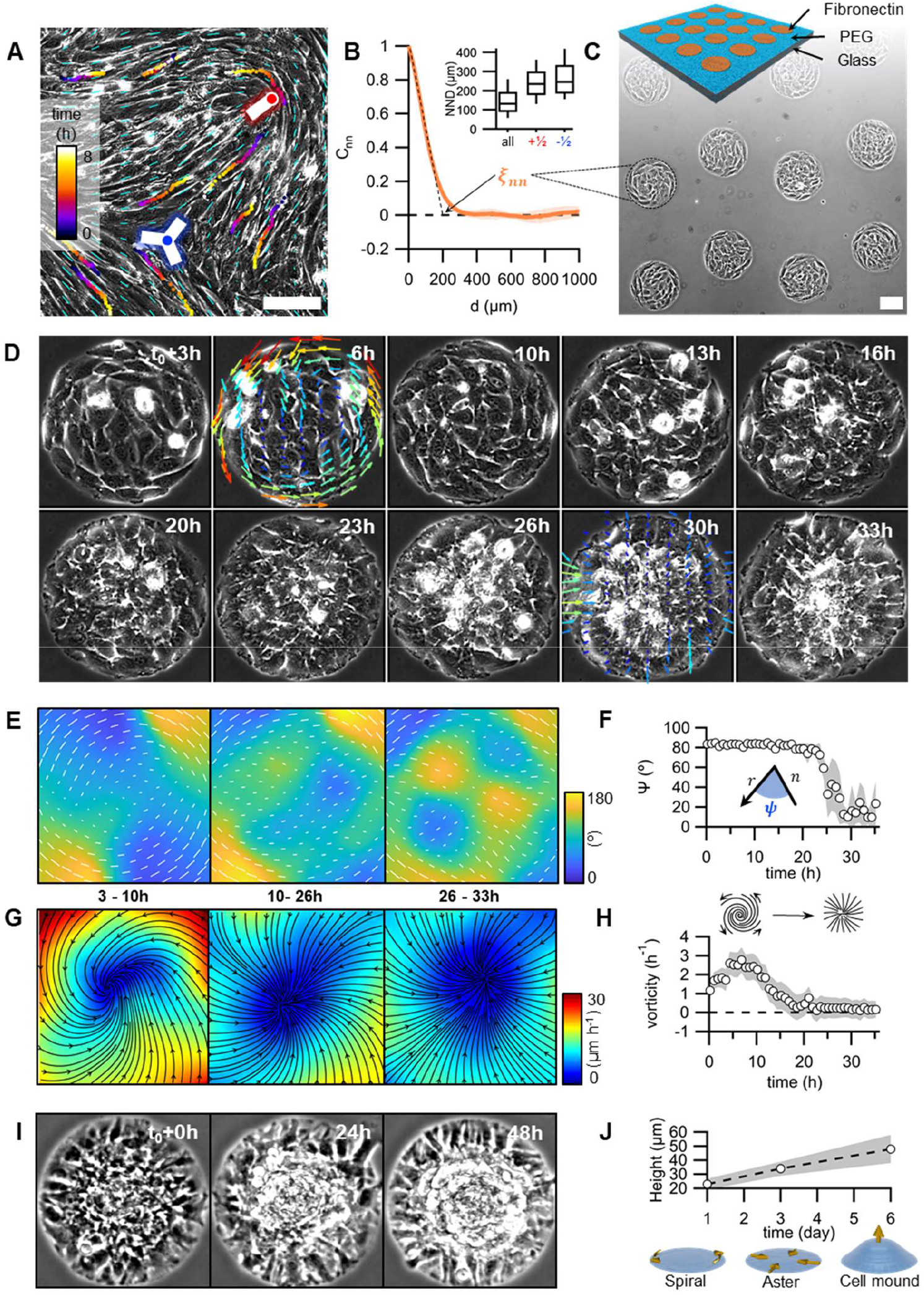
Myoblasts arrange into integer topological defects on circular micropatterns. (**A**) Phase contrast image of a confluent myoblasts monolayer. Cyan dashes indicate the orientational field. Positions of half-integer topological defects are shown (blue dot, s=−½; red dot, s=+½). Trajectories of single cells are depicted with color gradients. (**B**) Spatial autocorrelation function of the orientational field, *C*_*nn*_. *ξ*_*nn*_ is the nematic correlation length. Inset: nearest neighbor distance (NND) between topological defects. (**C**) Scheme of the micro-patterned surface and corresponding phase-contrast image with confined myoblasts (r=100μm). (**D**) Time series of a single myoblast disc (r=100μm). Time=0 at confluence. Velocity field is shown for 6 and 30h (v_max_=30μm/h). (**E**) Time-averaged orientational field calculated from D. Vectors and colormap depict local cellular orientation. Vector length corresponds to the local degree of order *S*. (**F**) Mean value of angle *ψ* between the local orientation and the radial direction over time (N=12). (**G**) Time-averaged flow field calculated from D. Flow directions are shown as black streamlines. The colormap depicts the local average velocity. (**H**) Average vorticity over time (N=12). (**I**) Time series of myoblast asters (r=100μm) forming mounds (**J**) Average height of myoblast assemblies with time. Scale bars, 100μm.

Inspired by previous studies(*25*–*29*), we reasoned that using circular confinement below the inter-defect characteristic length would induce cellular arrangements with one single defect with s=+1. Accordingly, C2C12 myoblasts were seeded on fibronectin-coated discs (Fig.1C) with diameters of the order of *ξ*_*nn*_. Reaching confluence on discs, C2C12 cells self-organized into spiral patterns, which exhibited persistent rotation for several hours (Fig.1D-H, 3-10h). Further proliferation led to the transformation of spirals into asters (Fig.1D-H, 10-26h), in which cells oriented radially from the center of the disc and ceased to rotate (Fig.1D-H, 26-33h, Fig.S2 and Movie S2). This transition from spirals to asters, theoretically predicted for active systems(*30*), correlated with an increase of total cell number (Fig.S3), suggesting that cell density controls it. Further proliferation led to the formation of cellular mounds at the centers of the asters (Fig.1I,J and Movie S3).

To investigate how cell mounds could emanate from integer defects we set out to study their structure and dynamics. First, we obtained average orientational and velocity fields from spirals stabilized by inhibiting proliferation with Mitomycin-C right after confluence (Fig.2A, Fig.S2, Movie S4 and Methods). The order parameter *S*, which measures the degree of orientational order, was minimal at the discs’ center and increased towards their boundaries (Fig.2B,C and Methods). The angle *ψ* between local orientation and the radial direction was 85±4° (N=12, Fig.S4 and Methods). Actin fibers in spirals, fluorescently-labelled with SiR-actin (Methods), showed a comparable *ψ* distribution (Fig. S5 and Movie S5). We used particle image velocimetry (PIV) to measure the velocity field within the cell monolayers (Methods). Near the boundaries, cell velocity was maximal at 27±7μm/h and, although it was dominantly azimuthal, as the direction of cell migration (Fig.2A), its radial component was non-vanishing (Fig.2B,C, N=12). The angle *β* between local orientation and velocity was thus not null, *β* =23±5° (Fig.S4, N=12), contrary to the general behavior of passive liquid crystals under shear(*5*). The role of active dynamics in the cellular arrangements was also evidenced by inhibiting myosin-driven contractility with Blebbistatin (Methods), which had a significant impact on the spirals’ shape. Although rotation persisted, *ψ* decreased to 55±5° (N=9), closer to the aster arrangement, and consequently, the radial velocity increased (Fig.S6).

**Figure 2.**
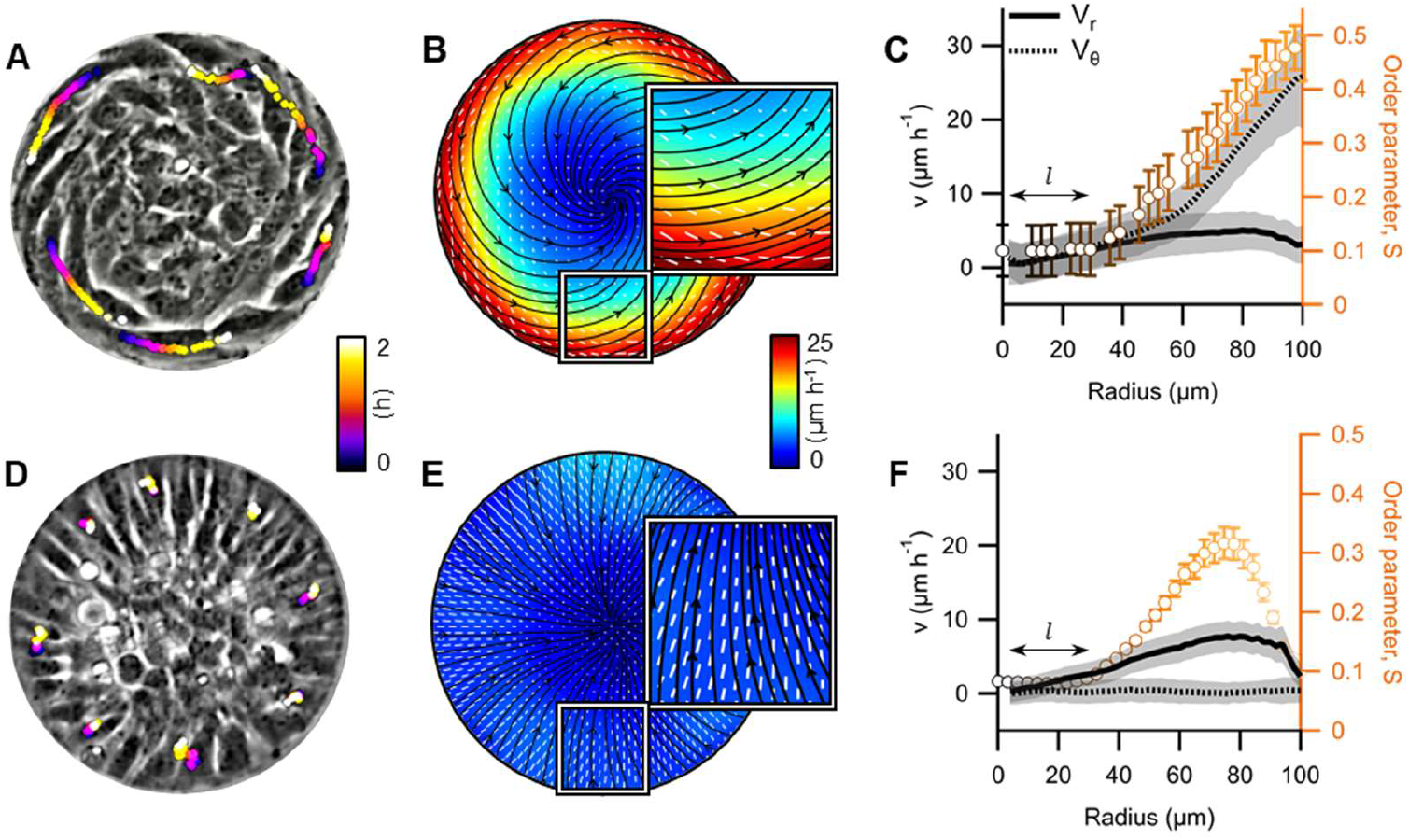
Cellular spiral and aster configurations. (**A**, **D**) Phase contrast images of spiral and aster arrangements, respectively. Colored dots depict the positions of some peripheral cells at different time points (r=100μm). (**B**, **E**) Average velocity and orientation fields (N=12 and 43, for spirals and asters, respectively). Streamlines indicate the direction of the cellular flow. Colormap represents average velocity. White vectors indicate local cellular orientation. Vectors’ length corresponds to the local degree of order *S* (**C**, **F**) Radial profiles of the azimuthal (v_θ_) and radial velocity (v_r_) components, and of *S*. *l* depicts the size of the defect’s core.

Above ~3·10^5^cells/cm^2^ (N=13), C2C12 cells formed stable asters (Fig.2D, Fig.S2 and Movie S5). Like in spirals, *S* increased towards the boundaries but both orientation and velocity were strictly radial (Fig.2D-F and Fig.S4). The radial velocity component (Fig.2E,F) was similar to that in spirals (Fig.2B,C). However, in asters, cells at the periphery remained almost immobile (Fig.2D and Movie S6). We thus suspected that the radial velocity originated from coherent actin flows. To show this, we fluorescently-labelled actin in asters with SiR-actin (Methods). PIV of fluorescence images revealed a radial velocity profile similar to the one observed by phase contrast microscopy (Fig.2E,F, Fig.S7 and Movie S7).

To elucidate the origin of the velocity and orientational patterns and infer their corresponding stress fields, we used a theoretical approach. Previous studies showed that bidimensional rotational flows can arise either from directional motion of dense active particles(*26*, *31*–*33*) or from gradients in anisotropic active stresses (*30*, *34*). We thus developed a 2D active nematic theory that accounted for both the directional motility of cells and the anisotropic active stresses. This theory is described elsewhere (*35*, *36*). To constrain the values of our parameters, we fitted our theoretical results to the experimental azimuthal velocity and orientational order profiles of spirals with radii of 50, 100, and 150μm (Fig.3A-C). Solutions leading to cell accumulation in the center of the asters, as observed experimentally (Fig.1I,J), comprised equal contributions of directional motility and active anisotropic cytoskeletal stress gradients(*35*, *36*). These solutions featured a compressive stress pattern that correlated with the cell density profile (Fig.3D-F).

**Figure 3.**
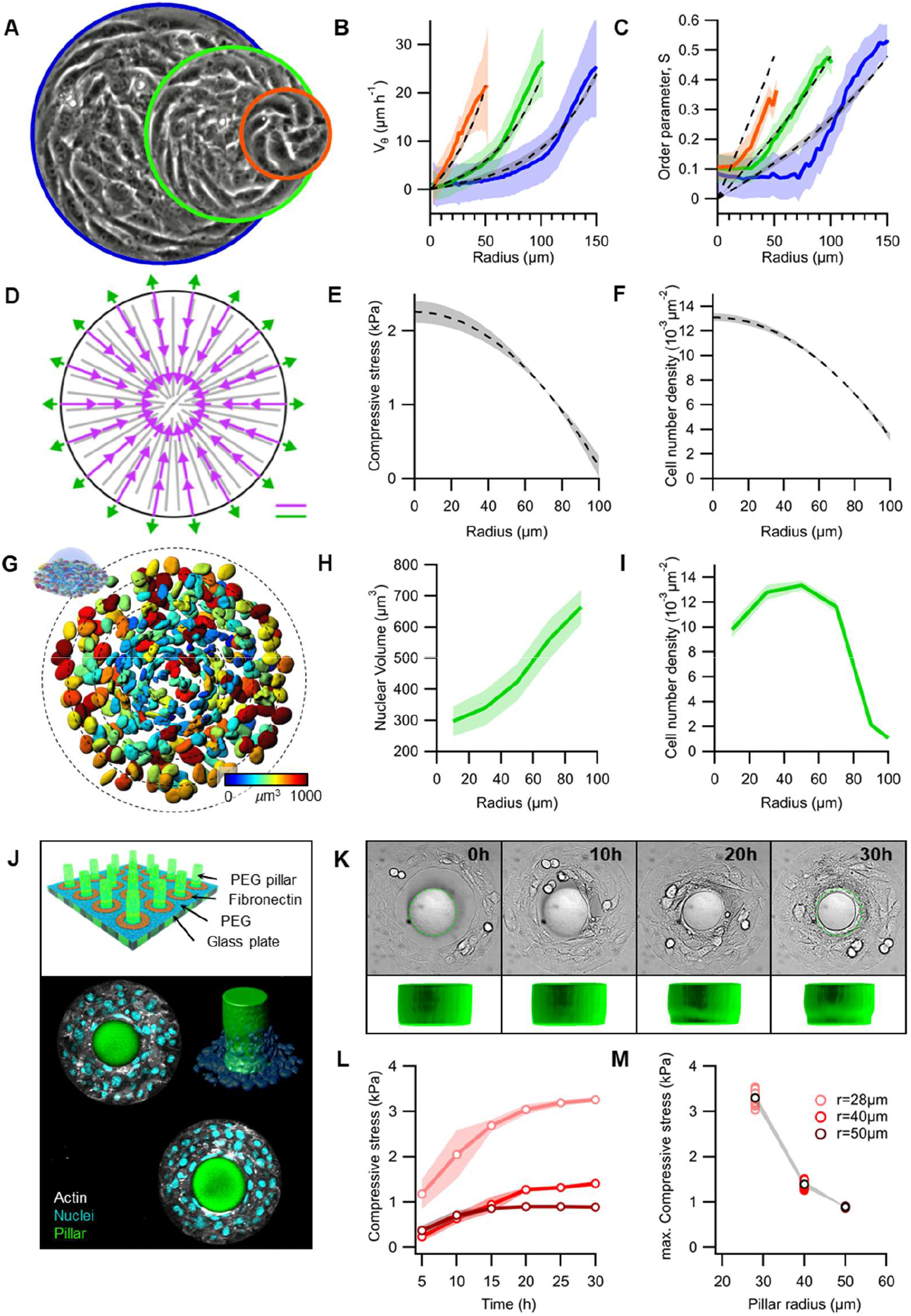
Integer topological defects concentrate active stresses. (**A**) Phase contrast images of spiral defects. Radii are 50, 100 and 150μm. (**B**) Average radial profiles of the azimuthal velocity (v_θ_) and (**C**) the order parameter S (N=11, 12 and 5 for r=50, 100 and 150μm, respectively). Theoretical fits are shown in dashed gray lines (solid magenta curves in Fig. 10 of (*36*)). (**D**) Steady-state active forces in asters: internal forces (purple) and forces at the boundary (green), see Sec. III of (*36*). (**E**) Steady-state compressive stresses and (**F**) cell density profiles in asters (solid magenta curves in Fig.11a and 11b of (*36*), respectively). (**G**) Nuclei at the base of a cellular mound (r=100μm). The colormap indicates the nuclear volume. (**H**) Radial profile of average nuclear volume and (**I**) average cellular density (N=10). (**J**) Scheme (top), fluorescence confocal composite and 3D rendering of the pillar constriction experiment (cellular ring, r_ext_=125μm). (**K**) Time-series showing the constriction of a pillar. Dashed line depicts the initial size of the pillar’s base (r=40μm). 3D rendering of the pillar is shown below. (**L**) Compression over time for pillars with different radii. (**M**) Maximum compression vs pillar radii. Scale bars, 100μm.

To test our theory (Fig3E,F), we characterized the nuclear volume and cell density, as internal rulers for external mechanical pressure(*37*). In asters, as expected, nuclear volume decreased towards the center (Fig.3G,H), and cell density increased (Fig.3G,I). To better assess the pressure at the core of the discs, we seeded myoblasts onto circular fibronectin rings enclosing non-adhesive elastic fluorescent micro-pillars with an elastic modulus E~4kPa (Fig.3J, Fig.S8,9 and Methods). Eventually, myoblast monolayers accommodated around the micro-pillars and compressed them (Fig.3K,L and Movie S8). The orientation and velocity fields were similar than for asters on discs, indicating that the pillars did not strongly affect the cellular arrangement (Fig.S10). We quantified the pressure around the pillars over time from their deformation (Fig.3L, Fig.S11 and Supplementary text). After ~30h, pressure plateaued at maximum values between 1-4kPa, being smaller for larger radii (Fig.3M). The good agreement with our theoretical results shows that the rotational flows in myoblast spirals result from an interplay between directional cell motion and anisotropic active stress gradients, both of which lead to compressive stresses in asters. We speculate that the pressure exerted at the core of asters is the promoter of cells’ displacement out of the monolayer’s plane and of the growth of cellular mounds.

To characterize the 3D organization of minimal cellular mounds (height~40μm), we fluorescently labelled actin and imaged z-stacks for several hours (Fig.4A and Methods). The 3D-averaged orientation and its azimuthal projection (Fig.4B and Supplementary text) revealed a peripheral layer of ordered cells, all through the z direction. Order was lost in the mounds’ center. To evaluate how the cell arrangement changed in the z direction, we extracted the radial distributions for the azimuthal angle *φ* and the zenith angle *ϕ* at different heights (Fig.4C,D). *φ* exhibited a bimodal distribution with peaks around 0 and 90°. Below ~10μm, *φ*~0° dominated, whereas above, *φ*~90° did (Fig.4C). Therefore, cells exhibited an aster arrangement at the bottom of the mound, and a spiral at the top. This aster-to-spiral transition was reverse of the transition in time discussed above (Fig.1D). We thus speculate that mound growth correlates with a decrease in cell density, which triggers this transition. Interestingly, *ϕ*~0° at the top of the mound, whereas an oblique component, *ϕ*~45°, appeared at the bottom (Fig.4D). This suggests that a vertical force, which could promote further growth of the mounds, results from the integration of multicellular stresses at the periphery.

**Figure 4.**
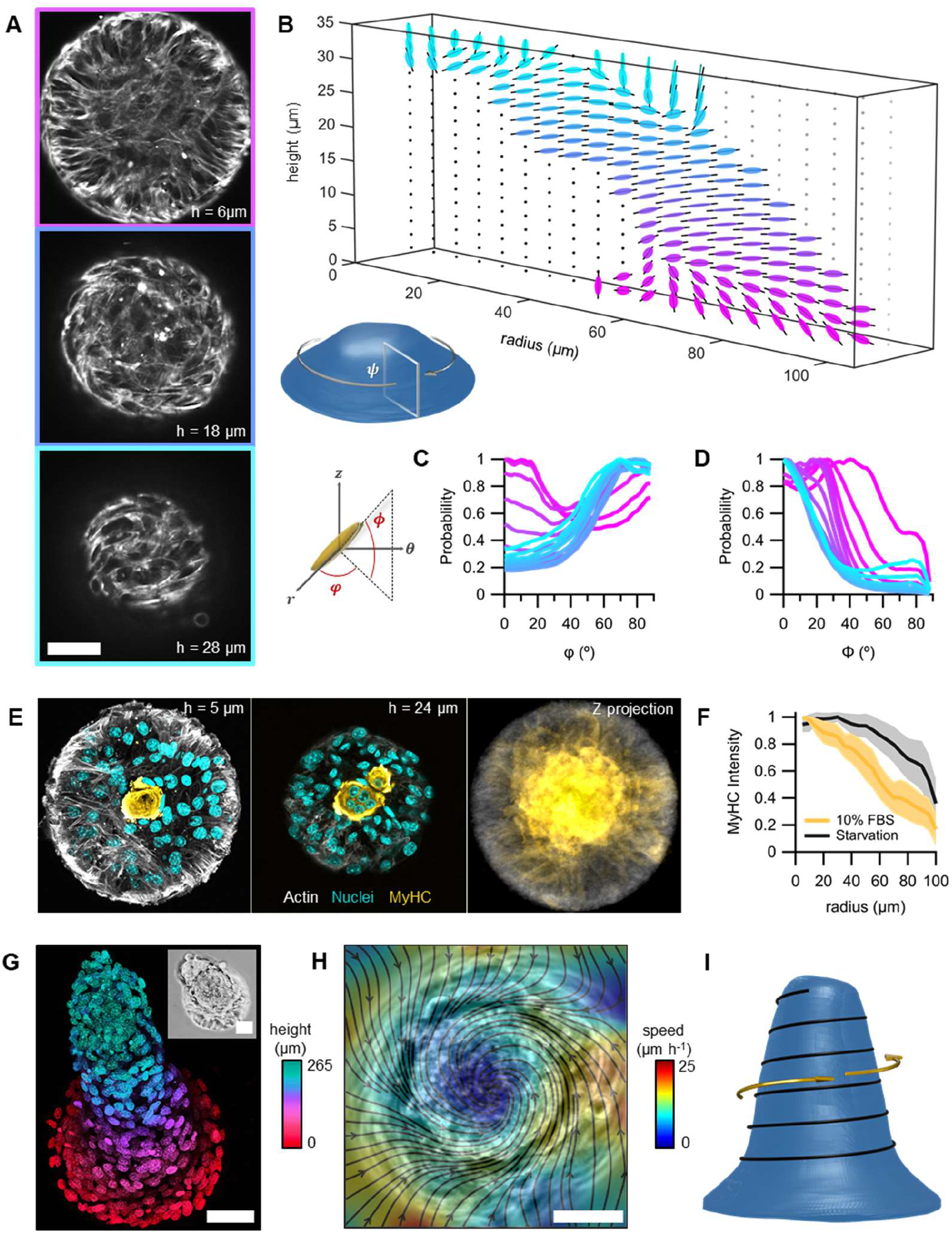
Topological defects organize 3D growth and differentiation. (**A**) Confocal sections of an actin-labelled cell mound. (**B**) Average section of the 3D orientational field. (**C**, **D**) Histograms of the angles *φ* and *ϕ*, respectively, for different heights. (**E**) Confocal sections showing the preferential position for myosin heavy chain (MyHC) expression (r=100μm). Left panel corresponds to the first layer of cells. Center panel corresponds to the midplane, displaying a multinucleated myotube-like structure. Right panel shows the maximum projection of MyHC intensity (N=43). (**F**) Radial profiles of averaged MyHC intensity under different conditions. (**G**) Z-projection of a nuclei-labelled cellular protrusion (r=100μm. Inset, phase contrast). (**H**) Average flow field from a cellular protrusion (r=100μm, phase contrast). (**I**) Proposed orientational field (black line) in 3D cellular nematic protrusions. Scale bars, 50μm.

Further evolution of myoblasts’ mounds depended on whether cells could differentiate or not. In the first case, myoblasts at the center of asters differentiated into globular multi-nucleated myotube-like structures after 6 days, as evidenced by the expression of myosin heavy chain (Fig.4E,F). In contrast, differentiation induced by serum deprivation led to canonical elongated myotubes with a much broader spatial centering (Fig.S12). Thus, we conclude that the stress patterns generated by cellular aster arrangements can trigger localized differentiation.

During morphogenesis, however, proliferating cells are usually not differentiated(*38*). Accordingly, we sought to inhibit differentiation in order to promote and study tissue morphogenesis. As previously observed (*39*), C2C12 cells cultured at high-passage numbers (>50) were unable to differentiate and featured higher proliferation rates. In this case, cellular mounds grew further in height, up to hundreds of microns (Fig.4G and Fig.S13), reminiscently of multicellular aggregation in amoeboid colonies(*40*). Considering that minimal cell mounds presented a spiraling top (FIG.4A-D), we imagine a similar arrangement could be preserved while growing into larger protrusions. Consistently, we observed collective rotational flows around the protrusion’s long axis (Fig.4H and Movie S9). These cellular structures were strictly dependent on confinement provided by the micropattern, as spontaneous degradation of the surrounding non-adhesive coating provoked their collapse (Movie S9). Thus, we conclude that minimal spiraling mounds (Fig.3A) can evolve into cylindrical vortices (Fig.4H,I), provided that their shape, growth and dynamics remain constrained by an integer topological defect.

In summary, our findings show how topological defects control the evolution of myoblast monolayers by localizing differentiation and steering morphogenesis. We foresee that topological defects could control multiple cell fate decisions and morphogenetic movements during development.

## Author contributions

P. G and A.R designed the research. P.G performed the experiments. C.B-M and K.K developed the theoretical model. All authors analyzed the data and participated in writing the manuscript.

## Data and materials availability

All data is available in the main text or the supplementary materials. High resolution movies, as well as the codes used for analyzing 3D orientational fields will be available upon reasonable request.

## Materials and Methods

### Cell culture and drug treatments

C2C12 mouse myoblasts were cultured in DMEM media containing 4500mg/L glucose, 1mM sodium pyruvate (Life Technologies) and supplemented with 10% Fetal bovine serum (FBS), 100units/mL penicillin and 100μg/mL streptomycin. Cells were maintained at 37°C under 5% CO_2_ and they were not allowed to become confluent. Maximum passages were kept below 20. For starvation conditions, used to induce differentiation, we supplemented the same DMEM media with 2% horse serum instead of 10% FBS.

For inhibiting differentiation, C2C12 cells were used after 50-60 passages, as previously reported(*39*). The cell batch was amplified and snap-frozen for future experiments.

For inhibiting proliferation, cells were treated with Mitomycin-C (Sigma) at 10μM for 1h at 37°C, then washed away and replaced by fresh medium. Imaging data were acquired up to 10h after treatment with Mitomycin-C, to avoid toxic effects(*41*).

For inhibiting contractility (myosin ATPase), cells were treated with Blebbistatin (Sigma) at 17μM.

In all treatments, DMSO concentration was kept below 10^−3^%v/v.

### Fluorescence labelling and imaging

For fluorescence immunostaining of myosin heavy chain, cells were fixed with 4% paraformaldehyde (Sigma) for 30min, permeabilized for 10min with 0.1% Saponin (Sigma) while blocked with 0.1% bovine serum albumin (BSA, Sigma). Finally, fixed cells are incubated 1h at room temperature with *Myosin* 4 Monoclonal Antibody conjugated with Alexa Fluor 488 (MF20, Thermofisher) at 5μg/mL (1:100 dilution) and 0.1% BSA.

Actin was labelled with SiR-Actin, conjugated with Alexa Fluor 647 (Spirochrome). Concentrations used were 1μM (30min incubation) for fixed samples and 250nM (6h incubation) for live imaging.

Cell nuclei were labelled after fixation with Hoechst 33342 (Thermofischer) at 10μg/mL (5min incubation).

Fixed samples were imaged by using a Nikon Eclipse Ti-E microscope equipped with a Nikon A1 confocal unit. We employed x40/x60 water/oil immersion objectives (NA 1.15/1.4). The microscope was operated with NIS-Elements software. A Zeiss LSM 710 upright confocal microscope (40x objective, NA 0.75) was used for imaging the top and exterior of cellular mounts. The microscope was operated with Leica Application Suite software.

### Time-lapse imaging

Time-lapse imaging was performed with an inverted microscope Nikon Ti-E installed into a thermostatically-controlled chamber (Life Imaging Technologies) and equipped with a micro-incubator for thermal, CO_2_ and humidity control (OKOlab). The microscope was also equipped with an automated stage and a Yokogawa CSU-W1 spinning disk unit. Image acquisition was performed with an Andor Zyla 4.2 Plus camera, operated with Slidebook Software. We performed fluorescence (60x lens, NA 1.4), phase contrast (10/20x objectives, NA 0.3/0.45) and differential interference contrast (DIC) imaging (20x lens, NA 0.45). 4D time-lapse was used for scanning actin-labelled cell mounds (60x lens, NA 1.4) and for the pillar compression experiments. The latter combined DIC and confocal fluorescence modes (20x lens, NA 0.45). Typically, we acquired 12 images/h for at least 10h.

### Substrate functionalization and micro-patterning

To prepare surfaces for micropatterning, glass bottom dishes (Mattek) were first activated using a plasma cleaner (Harrick Plasma, PDC-32G) for 3min. Then, the glass surface was treated with a 0.1mg/mL poly-lysine (PLL, Sigma) solution for 30min, then washed with HEPES buffer (pH=8.4). A solution of 50mg/mL mPEG (MW 5,000) - succinimidyl valerate (SVA) (Laysan Bio) was applied to passivate the surface for 1.5h, and then washed out with PBS. Substrates were normally used after preparation although they can be kept under PBS for 1-2 weeks at 4°C.

Micropatterns were generated by using a UV-activated mPEG-scission reaction, spatially controlled by the system PRIMO (Alvéole)(*42*), mounted on an inverted microscope Nikon Eclipse Ti-2. In the presence of a photo-initiator compound (PLPP, Alvéole), the antifouling properties of the PEGylated substrate are tuned by exposure to near-UV light (375nm). After illumination (1,200mJ/mm^2^) through a 20x objective PLL is exposed. After rinsing with PBS, fibronectin (Sigma) was incubated at 50μg/mL at room temperature for 5min in order to coat the PEG-free PLL motifs with the cell-adhesive protein. The excess of fibronectin was washed out with PBS. Patterned substrates were always used right after preparation. PBS was finally replaced by medium and a suspension of cells was added at densities of ~1·10^5^ cells/cm^2^. Samples were kept in an incubator at 37°C and 5% CO_2_. After 30min, non-adhered cells were washed out.

### Pillar constriction experiment

First, we fabricated mPEG-based fluorescent soft hydrogel micropillars. To this end, a “polymerization solution” was activated by near UV light with the system PRIMO (Alvéole), by inducing the photo-polymerization of pillars through illumination of our substrates with full circle motifs (see Fig. S8).

1. <u>Fluorescently-labelled Acrylate(AC)-mPEG:</u> Fluoresceinamine (Sigma) was dissolved at 0.1mg/mL in HEPES buffer solution (pH=8.3) and mixed with an equal volume of a 50mg/mL AC-PEG (MW 2,000)-SVA (Laysan Bio) solution, which was prepared in the same buffer. The resulting mixture was vortexed and let sit at room temperature in the dark for 1h.
2. <u>Polymerization mixture:</u> to prepare 100 μL of a 5% PEG hydrogel we mixed 50μL of a 10% solution of 4arm-PEG (MW 20,000)-AC (Laysan Bio) in water, 25μL of fluorescently-labelled AC-mPEG, 24μL of water and 1μL of 3- (Trimethoxysilyl)propyl methacrylate (Sigma). The water employed was double-distilled and it was degassed by flowing Argon for 5min.
3. <u>Substrate preparation:</u> a glass bottom dish (Mattek) was first activated with a plasma cleaner (Harrick Plasma, PDC-32G) for 3 min and a PDMS stencil (thickness, 300 μm, Alvéole) with 4 small circular wells (r=2mm) was placed onto the glass substrate. 30-50μL of the polymerization mixture was added into each of the circular wells, which were then covered by a polyethylene film, 100μm thick. Micro-pillars were fabricated right after.
4. <u>Fabrication of micro-pillars:</u> the PRIMO system (Alvéole) was used to fabricate micro-pillars ~300μm high. To this end, we illuminated the upper base of the glass substrates with a 10x objective. Full circle motifs (r=75μm) of UV-light (100-200mJ/mm^2^) were projected. After photo-polymerization, the remaining solution was washed out and rinsed with PBS.
5. <u>Micro-patterning:</u> we functionalized the glass substrate with PEG and generated ring micro-patterns around the micropillars by using the same protocols above (see Section “Substrate functionalization and micro-patterning). Here, ring motifs (r_int_=75μm, r_ext_=125μm) were manually located with the software Leonardo (Alvéole). Subsequently, substrates were incubated with fibronectin and finally, cells were seeded, also as explained above.

### Fabrication of PEG hydrogel disks and measurement of their elastic properties

In order to characterize the elastic modulus of the PEG hydrogels we prepared 300μm films of 4arm-PEG (MW 20,000) with different densities (20, 10, 5 and 2.5%w/v). The protocol employed for the preparation of the “polymerization mixture” was the same as the one above (see Section “Pillar constriction experiment”). However, in this case we added our “polymerization solution” in the circular wells (14mm diameter, 1mm high) of glass-bottom dishes (Mattek). We covered the well with a glass coverslip to ensure obtaining a flat surface. Subsequently, we polymerized the whole content of the wells by illuminating the samples 1min in a UV-curing chamber (375nm, Asiga Flash), obtaining hydrogel disks 14mm in diameter and 1mm thick. After polymerization, gels were rinsed with double-distilled water kept wet at 4°C.

Force-displacement curves were obtained by using a FT-S100 micro-force sensing spherical probe (r=250μm, Femtotools). We performed 9 indentations for each gel at 2μm/s and obtained Force vs Displacement curves (Fig S9, A, B). Calibration hydrogels (4 and 50kPa, Petrisoft) were employed to complete the measurements. The elastic modulus of the lowest density gel (2.5%) was assumed to be 4kPa (Fig. S9, C).

### Image analysis

#### Flow, orientation and associated quantities

Tracer-free velocimetry analysis of the flows in the cell monolayers was performed with a public domain particle image velocimetry (PIV) program implemented as an ImageJ plugin(*43*). Manual Tracking ImageJ plugin was used to manually track trajectories of cells. The cell shape 2D orientational field **n** was extracted by using the imageJ plugin OrientationJ, which is based on the structure tensor method(*44*). The angle *θ* is the local orientation of **n** with respect to a fixed axis. The amplitude of **n**, named coherency C, was also extracted from the imageJ plugin OrientationJ. 3D cell-shape orientation analysis was based on the same method but considering intensity gradients in 3D (see Supplementary text, below). It was implemented as a MatLab function. Further analysis from velocimetry and orientation data were also performed with custom-written Matlab codes.

We used the average vorticity to assess the rotational component of the flow field. Local vorticity was calculated as the curl of the (2D) velocity vector field obtained from PIV:

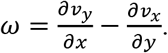

The spatial nematic autocorrelation function *C*_*nn*_ (Fig.1B and Fig. S1C) was calculated from each orientational field position like

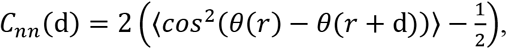

where *θ* is the local orientation of **n** with respect to a fixed axis. We considered *C*_*nn*_ at timepoints within one-hour period for temporal averaging. The characteristic nematic length *ξ*_*nn*_ was extracted from the intersection of the initial linear decay and *C*_*nn*_=0.

The angle *ψ* was obtained for each position from the scalar product between the orientation vectors **n** and their corresponding radial direction vectors 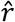. The angle *β* was obtained for each position from the scalar product between the orientation vectors **n** and their corresponding velocity vectors 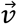. Note that both angles *ψ* and *β* were considered only if the coherence of their corresponding orientation vectors(*44*) presented values superior than a certain coherence threshold value, which is specified in each figure caption. Mean values for *ψ* and *β* were obtained from the Gaussian fits of their corresponding probability distributions.

To obtain the orientational order parameter *S*, we first computed the nematic order tensor Q from the orientation field **n**. Specifically, the components of the nematic tensor were

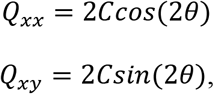

where C corresponds to the coherency and *θ* is the local orientation of **n** with respect to a fixed axis. The nematic order parameter *S* was calculated from the time-averaged components of the nematic tensor *Q* for each position like

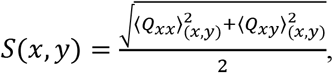

where the brackets 〈 〉 denote a time average over a local position in the space matrix.

#### Nearest neighbor distance between half integer defects

For the detection of half-integer defects for the nearest neighbor analysis in Fig. 1B we build up on previously used algorithms. First, we define as defect areas, the regions where the parameter 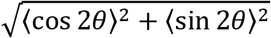 was below a threshold value. The brackets 〈 〉 denote an average over a local region. In order to assess their topological strength, we calculated the winding number 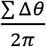, where ∑ Δ*θ* is the accumulated rotation of the orientational field around these low-order regions (*45*). Finally, for the nearest neighbor distance analysis, we compared all the distances between defects and selected the minimum values corresponding to each pair.

#### Volume segmentation of nuclei and pillars

Segmentation of nuclei and pillars, as well as the quantification of their volume were performed with Imaris software (Oxford Instruments). Nuclear volume and nuclear density profiles were calculated with custom-written Matlab codes.

#### Calculation of errors

In general, error bars correspond to the standard error of the mean (SE), calculated like 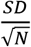, where SD is the standard deviation obtained considering all timepoints per experiment, including different replicates. N corresponds to the number of replicates. Thus, note that we did not use temporal averaging for the calculation of the errors.

Errors associated to the mean values of the angles *ψ* and *β* were calculated as 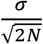, where *σ* is the width (SD) of the gaussian function 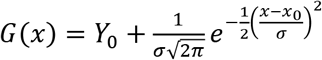 fitted to the angles’ probability distributions.

## Supplementary text

### 1. Linear elastic cylinder subjected to a uniform pressure

In the following, we derive the theoretical equation used to quantify the cellular forces exerted on deformable elastic pillars.

The geometry of pillars is approximated by a cylinder of radius *R* and height *h*. Due to the symmetries of the cylinder, we focus on axisymmetric solutions, (i.e. independent on the azimuthal coordinate *θ*) and use cylindrical coordinates (*r*, *θ*, *z*). Furthermore, we consider that our deformable elastic pillars behave as a linear elastic material (Fig. S9, A), so that the stress tensor *σ* obeys

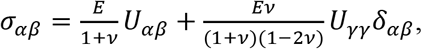

where the symmetric part of the strain tensor is 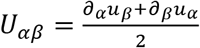 with **u** being the displacement vector. The material parameters are the elastic modulus *E* and the poisson ratio *v*.

The force balance equation reads

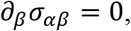

as there are no bulk forces applied on the pillars.

To complete our description, we need to specify the boundary conditions. On the lateral surface of the cylinder, we consider that cells exert a uniform compressional stress −*P*, so that *σ*_*rr*_(*r* = *R*) = −*P* and *σ*_*zr*_(*r* = *R*) = 0. On the bottom surface of the cylinder, we consider vanishing displacement in the z-direction *u*_z_(*z* = 0) = 0. On the bottom surface, the cylinder is allowed to slide freely in the radial direction, so that *σ*_*zr*_(*z* = *h*) = 0. On the upper surface of the cylinder, we consider stress free conditions *σ*_*zz*_(*z* = *h*) = 0 and *σ*_*rz*_(*z* = *h*) = 0.

A steady-state solution to the above problem, corresponds to a displacement field **u** with 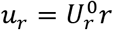, *u*_*θ*_ = 0, and 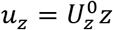. In this case, the non-vanishing components of the stress tensor are:

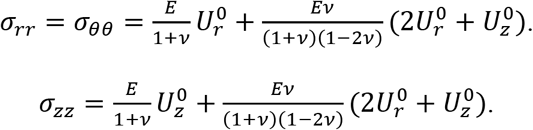

By enforcing that *σ*_*zz*_(*z* = *h*) = 0 and *σ*_*rr*_(*r* = *R*) = −*P*, we obtain the displacement field

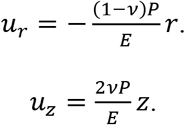

Rewritten in terms of the areal strain, 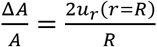, reads

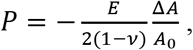

where Δ*A* is the difference of cylinder base’ areas after deformation and *A*_0_ is the area of the cylinder’s base before deformation.

This equation was used to quantify the cellular forces exerted on deformable elastic pillars. The areal strain was calculated from the experimentally-measured pillars’ volume like

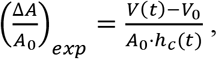

where *V* corresponds to the segmented volume of the cylindrical pillar at a fixed height, typically 40μm, which was larger than the compressed height *h*_*c*_. Temporal evolution of *h*_*c*_ was assesed manually from 3D volume renderings of the pillars (Fig. S11). *V*_0_ and *A*_0_ correspond to the initial segmented volume and base area of the cylindrical pillar, respectively. The elastic modulus *E* was extracted from indentation measurements (Fig. S9). For the calculation of forces on the PEG pillars we considered PEG hydrogels to be incompressible, thus *v* = 0.5(*46*).

### 2. 3D director field from fluorescence confocal z-stacks

In the following, we explain the procedure to determine 3D director field **n** from z-stacks of fluorescence images.

Let us consider a 3D intensity map *I*(*x*, *y*, *z*), such as a z-stack of fluorescence images, where (*x*, *y*, *z*) represents the cartesian coordinates. First, using the Matlab function *interp3*, we interpolated the 3D intensity map *I*(*x*, *y*, *z*) so that the resolution of the *z* coordinate matched the resolution of the (*x*, *y*)-planes. Next, we used the Matlab function *imgaussfilt3* to apply a Gaussian filter with standard deviation σ_1_ on *I*(*x*, *y*, *z*). This part eliminated small-wavelength fluctuations from the intensity map. For each pixel, we computed the gradient of the intensity map (*∂*_*x*_*I*, *∂*_*y*_*I*, *∂*_*z*_*I*) by using the Matlab function *imgradientxyz*. We computed the structure matrix, which is defined as

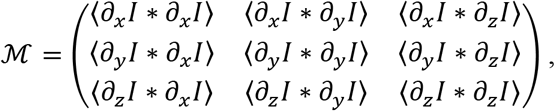

where the brackets 〈·〉 denote a second Gaussian filter with standard deviation σ_2_. We defined the traceless structure matrix as 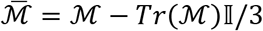, where *Tr* denotes the trace operator and 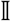 denotes the identity matrix. For each pixel, the matrix 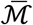 was diagonalized by the Matlab function *eig*. For each pixel, the three eigenvectors of 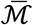 define, in general, an orthogonal basis. The eigenvector with the smallest eigenvalue represented the direction of smallest variation of the intensity map *I*(*x*, *y*, *z*) in the vicinity of the pixel (*x*, *y*, *z*). We considered the director field **n** parallel to the eigenvector with minimal eigenvalue. Note that, the orientation of **n** was determined up to a sign, meaning that **n** → −**n** were indistinguishable. We choose *n*_*z*_ to be positive. The amplitude of **n** was set by the smallest eigenvalue of 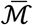. For each pixel, we computed the components of the nematic tensor field Q in cylindrical coordinates, taking as the center the geometrical center of the confining domains. Finally, we averaged the components of the nematic tensor field Q over time and experiments. In conclusion, the method presented two input parameters given by the standard deviations of two Gaussian filters, and outputted a nematic tensor field Q from a 3D intensity map *I*(*x*, *y*, *z*).

To construct Fig. 4B, we apply the above-mentioned routine with the input parameters σ_1_ = 1 px and σ_2_ = 5 px to actin-stained cell mounds (N=8) and obtained the averaged nematic tensor field Q in cylindrical coordinates. We binned the data in the radial direction so that 20 points are shown. To represent the data, we used the following procedure. First, for each data point, we computed the eigenvectors and eigenvalues of the binned nematic tensor field using *eig*. Next, for each data point, we constructed a 3D ellipsoid of revolution with the major axis proportional to the largest eigenvalue and the minor axes proportional to the mean of the two lowest eigenvalues. For each z-plane in Fig. 4B, only the 3D ellipsoids of revolution that had a trace of the binned nematic tensor larger than the mean of each plane, were shown.

## Supplementary figures

**Figure S1.**
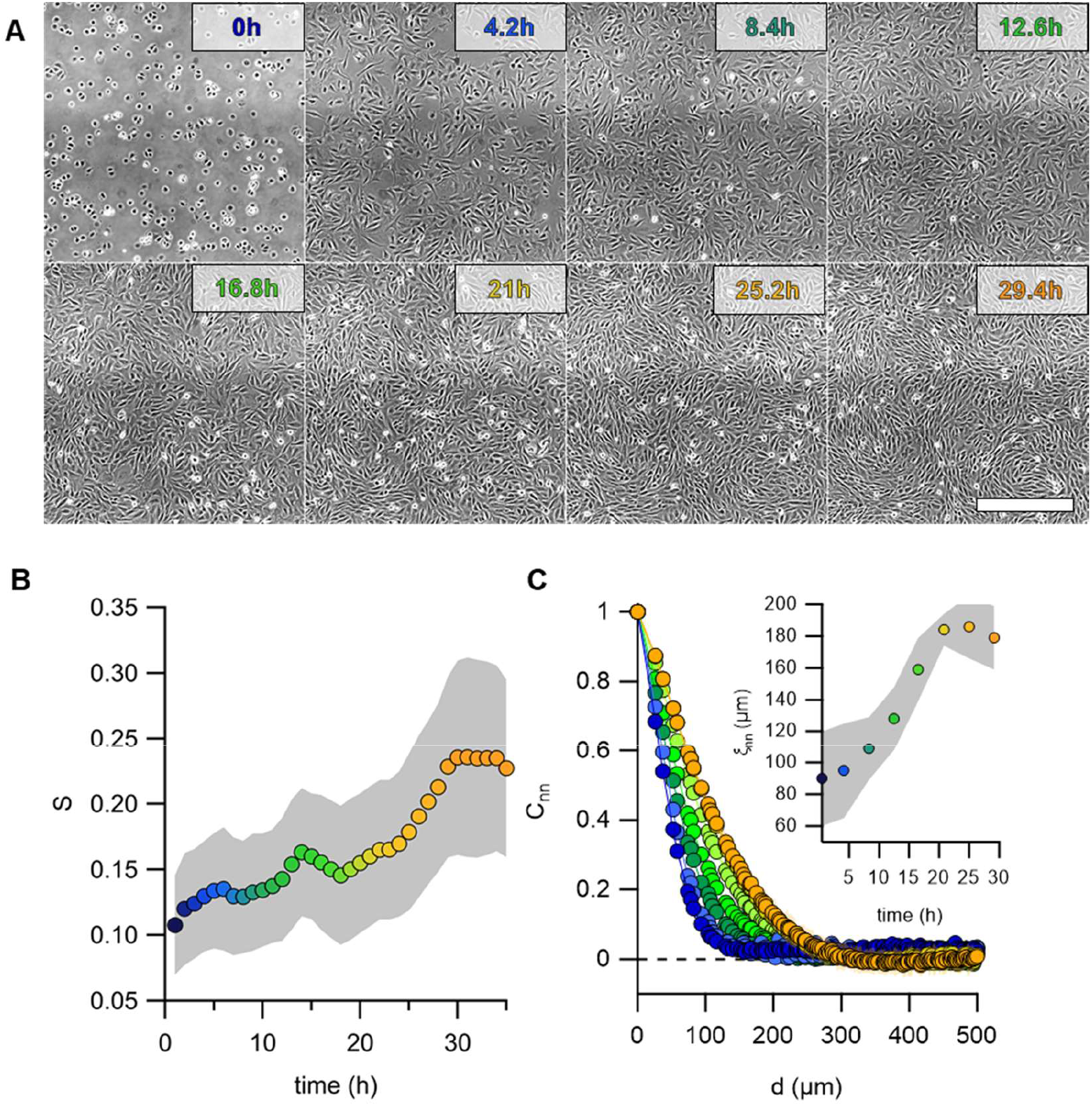
Formation of a nematic cellular monolayer. (**A**) Time series of a proliferating monolayer of C2C12 myocytes. Scale bar, 500μm. (**B**) Average order parameter S in function of time. (**C**) Temporal evolution of the spatial autocorrelation function C_nn_ and nematic autocorrelation length **ξ**_nn_ (inset).

**Figure S2.**
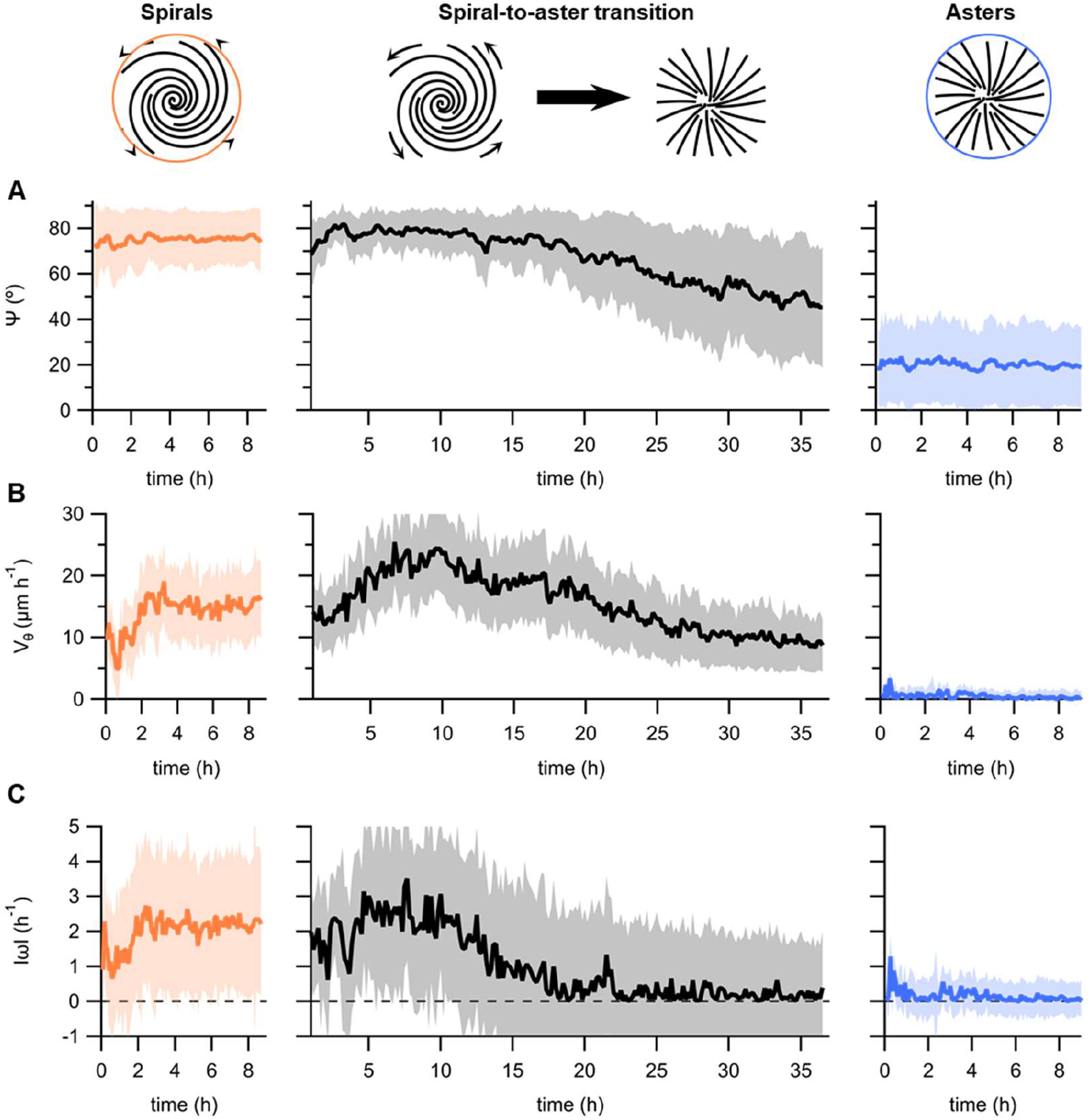
Morphology and dynamics for spirals and asters. Temporal evolution of **(A)** *ψ*, (**B**) azimuthal velocity (v_θ_) and (**C**) vorticity *ω*, in spirals (orange, N=12), during the spiral-to-aster transition (black, N=12) and in asters (blue, N=43).

**Figure S3.**
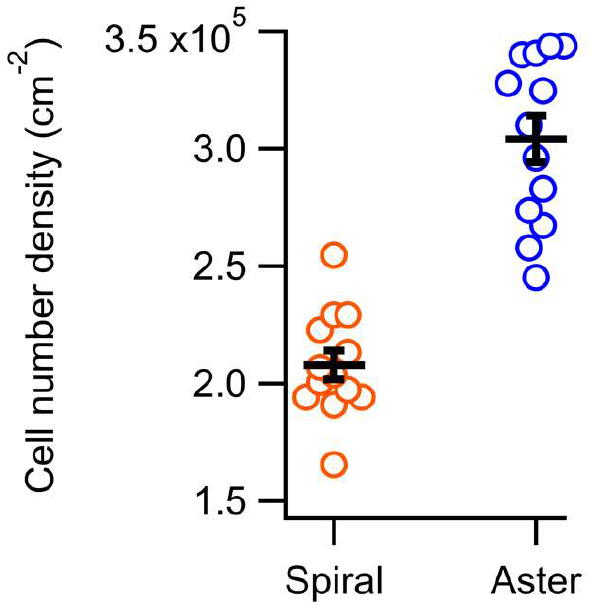
Cell number density in spirals and asters. N=13 both for spirals and asters.

**Figure S4.**
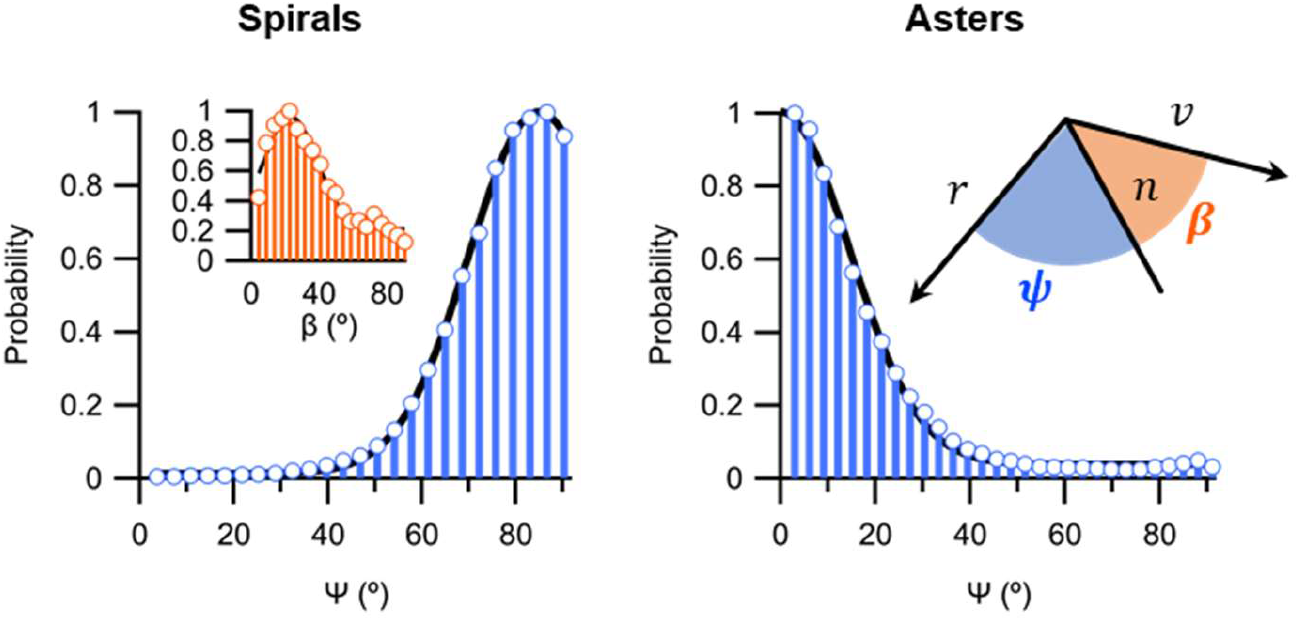
Distribution of angles *ψ* and *β* and for spirals and asters. *ψ* values (blue bars) are calculated from orientation vectors at distances r<0.9R and r>0.6R, being R the radius of the islands, and with S>0.3. For spirals, *β* values (orange bars) are calculated from all the vectors from the average velocity and orientational vector fields (Fig. 2B and E). N=12 and 43, for spirals and asters, respectively. Black lines correspond to Gaussian fits.

**Figure S5.**
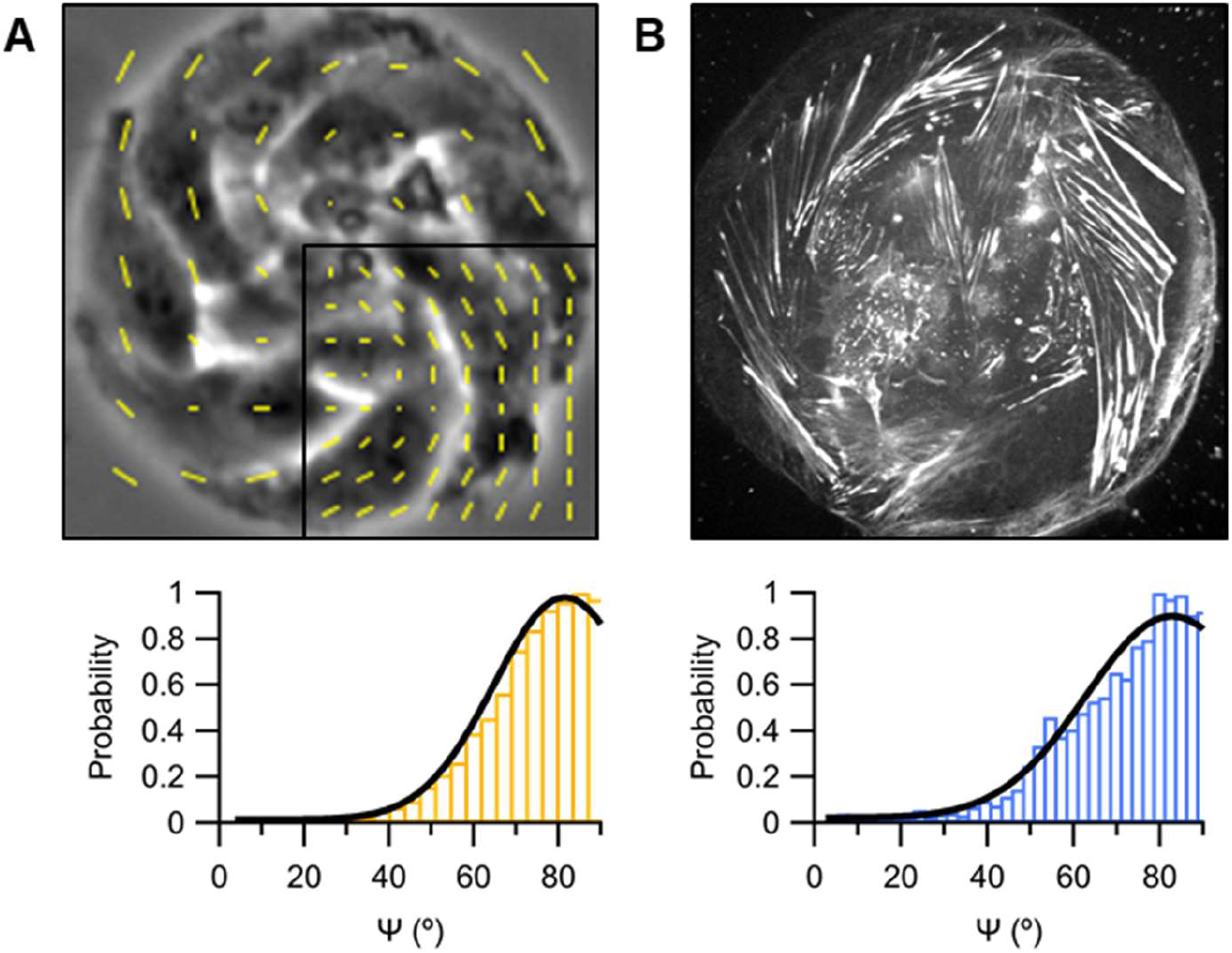
Orientational field from cell-shape and actin fibers. (**A**) Phase contrast image of a stabilized spiral (r=50μm). Yellow vectors correspond to the local orientation. For clarity, all extracted vectors are only shown in the framed inside panel. (**B**) Confocal micrograph of the bottom plane of an F-actin-labelled stabilized spiral (r=50μm). Bottom histograms show the distribution of the values for the angle *ψ* extracted from time-lapses represented by images in A (N=11) and B (N=7). Values considered correspond to vectors at distances r<0.95R and r>0.3R, being R the radius of the islands, and with S>0.4.

**Figure S6.**
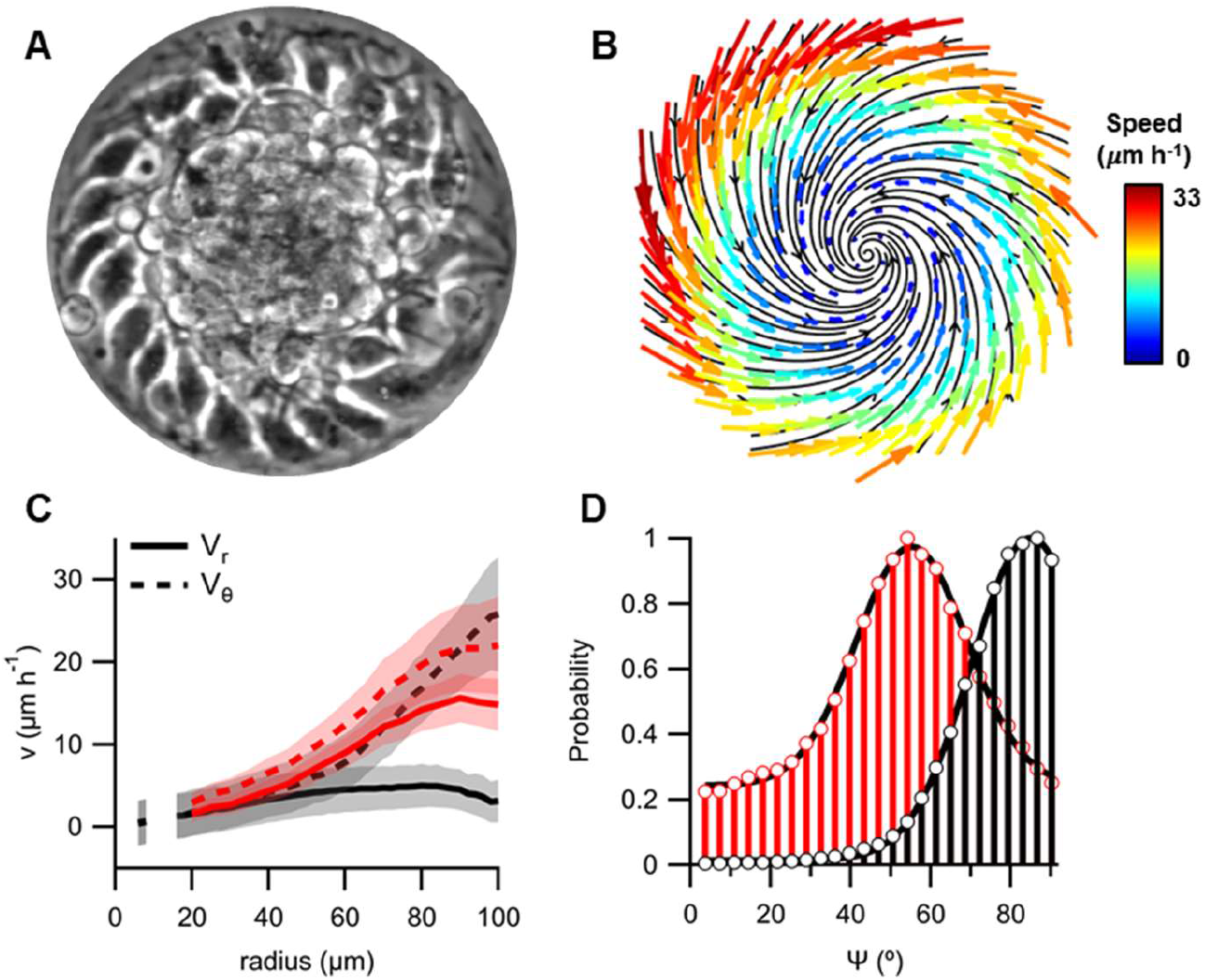
Influence of myosin activity on spiral’s morphology and dynamics. (**A**) Phase contrast micrograph of a Blebbistatin-treated spiral (r=100μm). (**B**) Average flow field (N=9). (**C**) Radial profiles of the radial and azimuthal velocity components for spirals, treated (red) or not treated (black) with Blebbistatin. (**C**) Radial profiles of the radial and tangential velocity components. (**D**) *ψ* values for spirals, treated (red) or not treated (black) with Blebbistatin. Values considered correspond to vectors at distances r<0.9R and r>0.6R, being R the radius of the islands, and with S>0.3.

**Figure S7.**
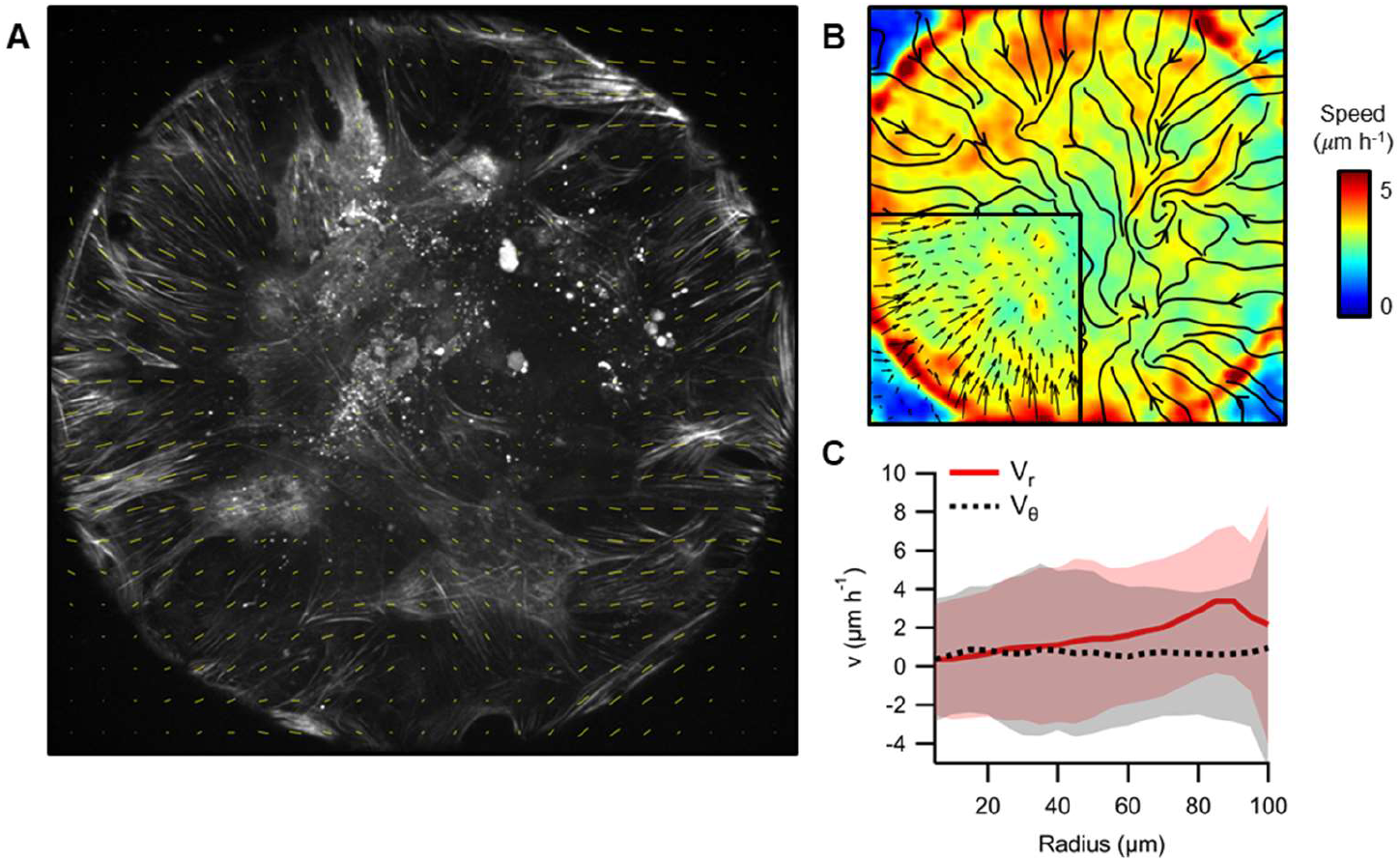
Actin flows in asters. (**A**) Confocal micrograph of an aster base (r=100μm). Filamentous actin was labelled with SiR-Actin. Yellow lines indicate the local average orientation. (**B**) Average flow field (N=4). Streamlines and vectors (inset) indicate the direction of the actin flow. The colormap indicates the average speed. (**C**) Radial profiles of the radial (red) and azimuthal (black) velocity components.

**Figure S8.**
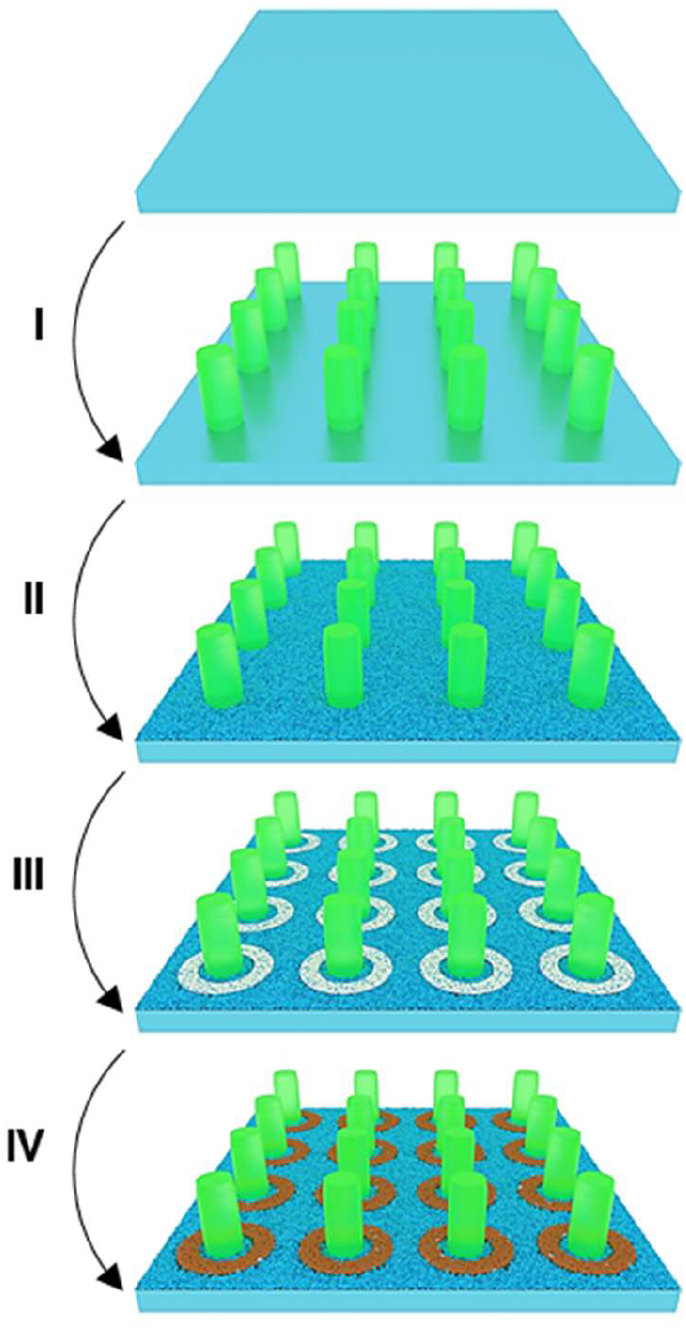
Micro-pillar compression experiment. Schematics of the protocol employed to fabricate cell-adhesive rings enclosing passive fluorescent hydrogel micro-pillars. After activation of the glass substrate, micro-pillars were fabricated by illuminating a photo-polymerizable mPEG solution with disk patterns of UV light with an inverted microscope (step I). Then, the substrate was functionalized with PLL-PEG (step II). PEG chains were locally photo-degraded by illuminating the substrate with ring patterns of UV light (step III). Finally, fibronectin was incubated (step IV).

**Figure S9.**
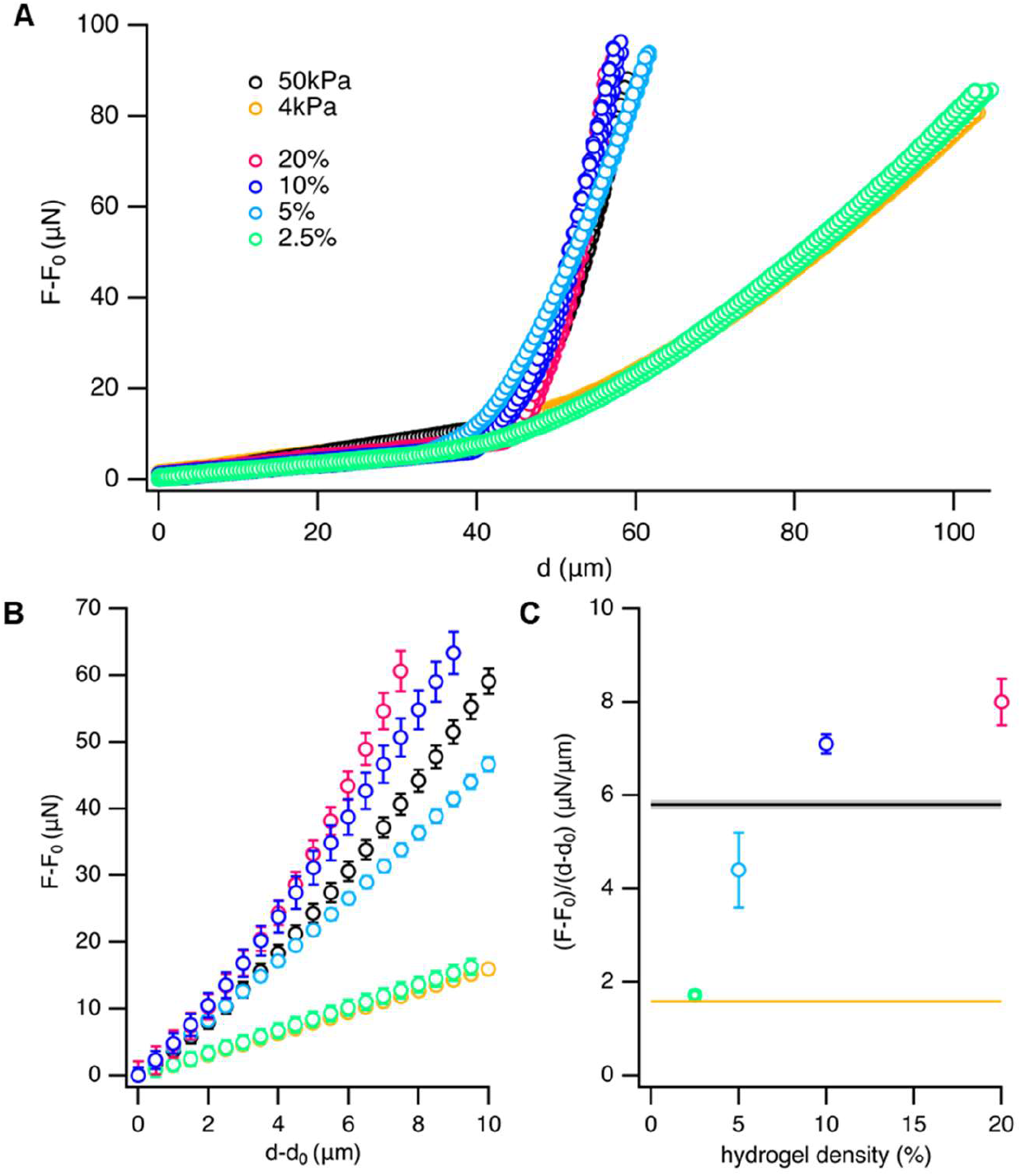
Elastic modulus of hydrogel films. (**A**) Force-displacement curves (N=9) for gels of different densities and calibration gels with E=4 and 50kPa. (**B**) Averaged curves from A. Data is normalized by the distance at which indentation starts (d0, inflection points in A). (**C**) Average slopes from curves in B prior averaging (N=9).

**Figure S10.**
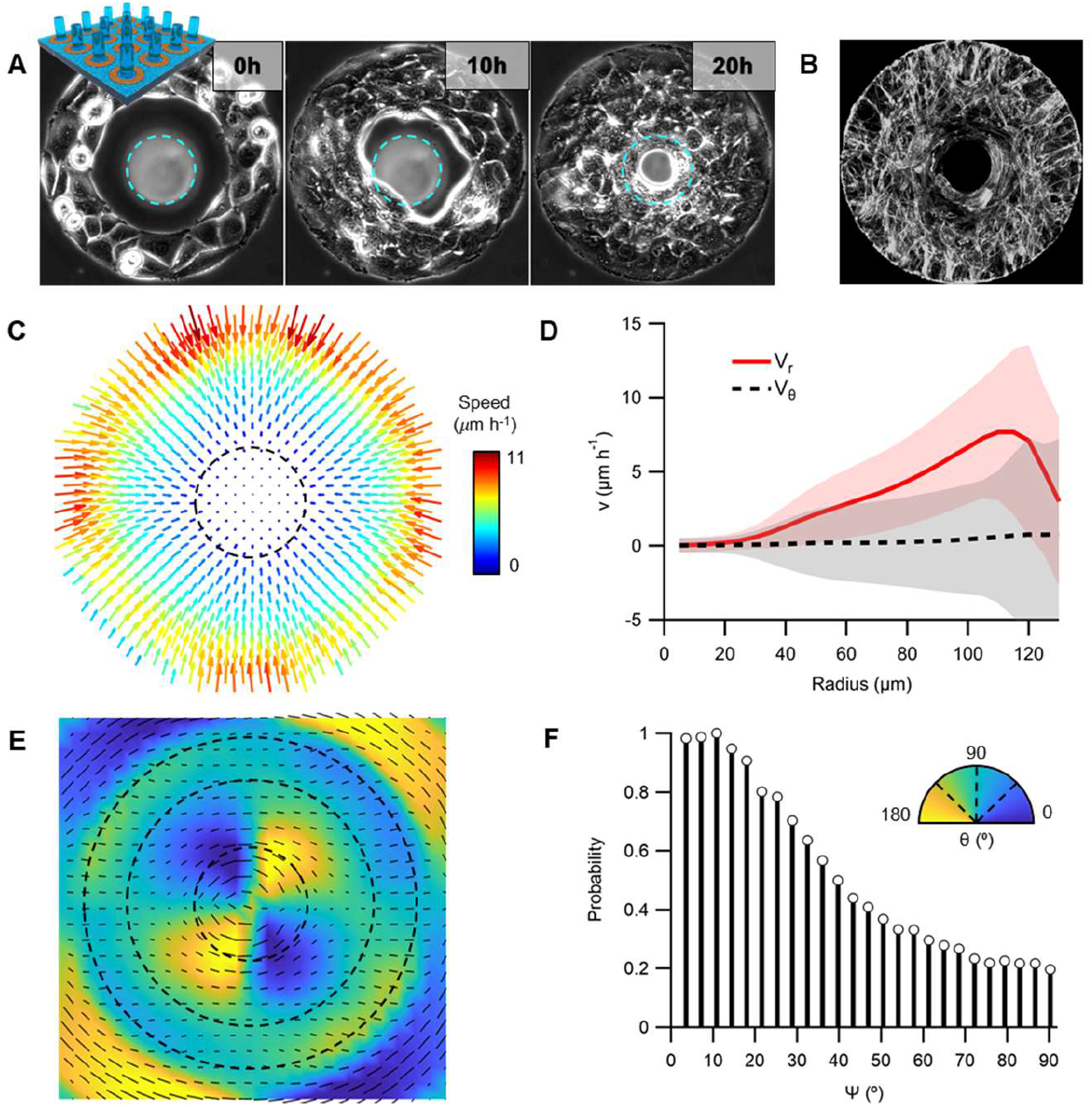
Formation of aster arrangements around pillars. (**A**) Time-series of C2C12 cells constricting a hydrogel micro-pillar. Cellular rings, r_ext_=125μm. Cyan dashed line indicates the initial pillar’s section. (**B**) Actin-stained cells after constriction show an aster arrangement. (**C**) Average flow field of flows around pillars (N=9). (**D**) Average radial profiles of the radial (red) and azimuthal (black) velocity profiles. (**E**) Average orientational field. For clarity, only half of the total number of orientation vectors are shown. (**F**) *ψ* distribution. *ψ* values are calculated from vectors at distances r<0.85R and r>0.65R, being R the radius of the islands, and with S>0.3.

**Figure S11.**
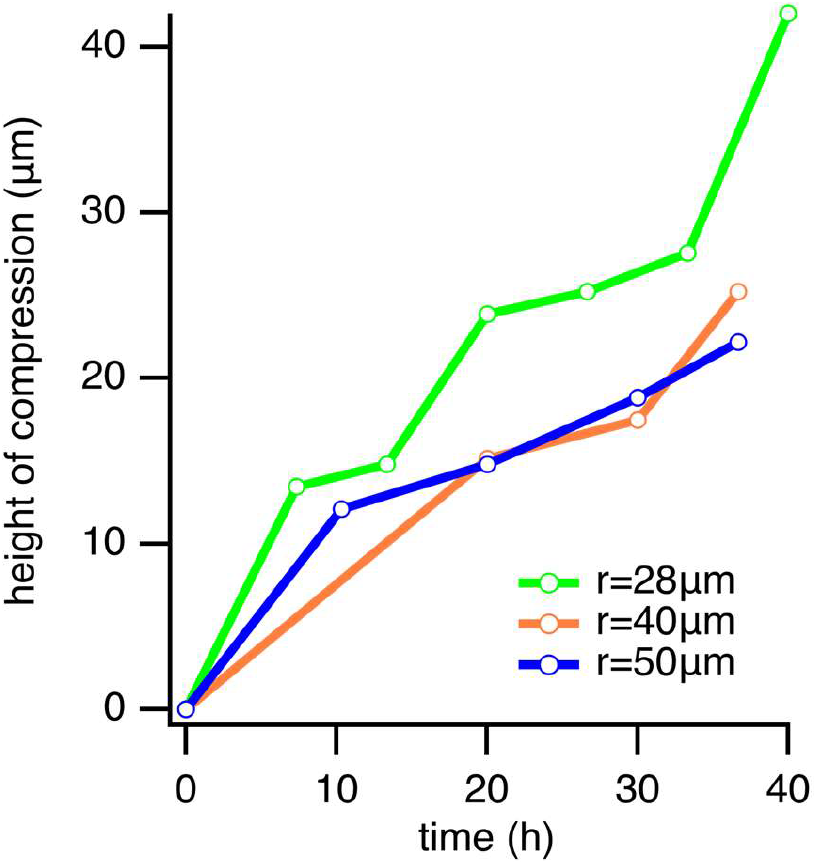
Height of pillars’ compression. Temporal evolution of compression’ height for pillars with different radii.

**Figure S12.**
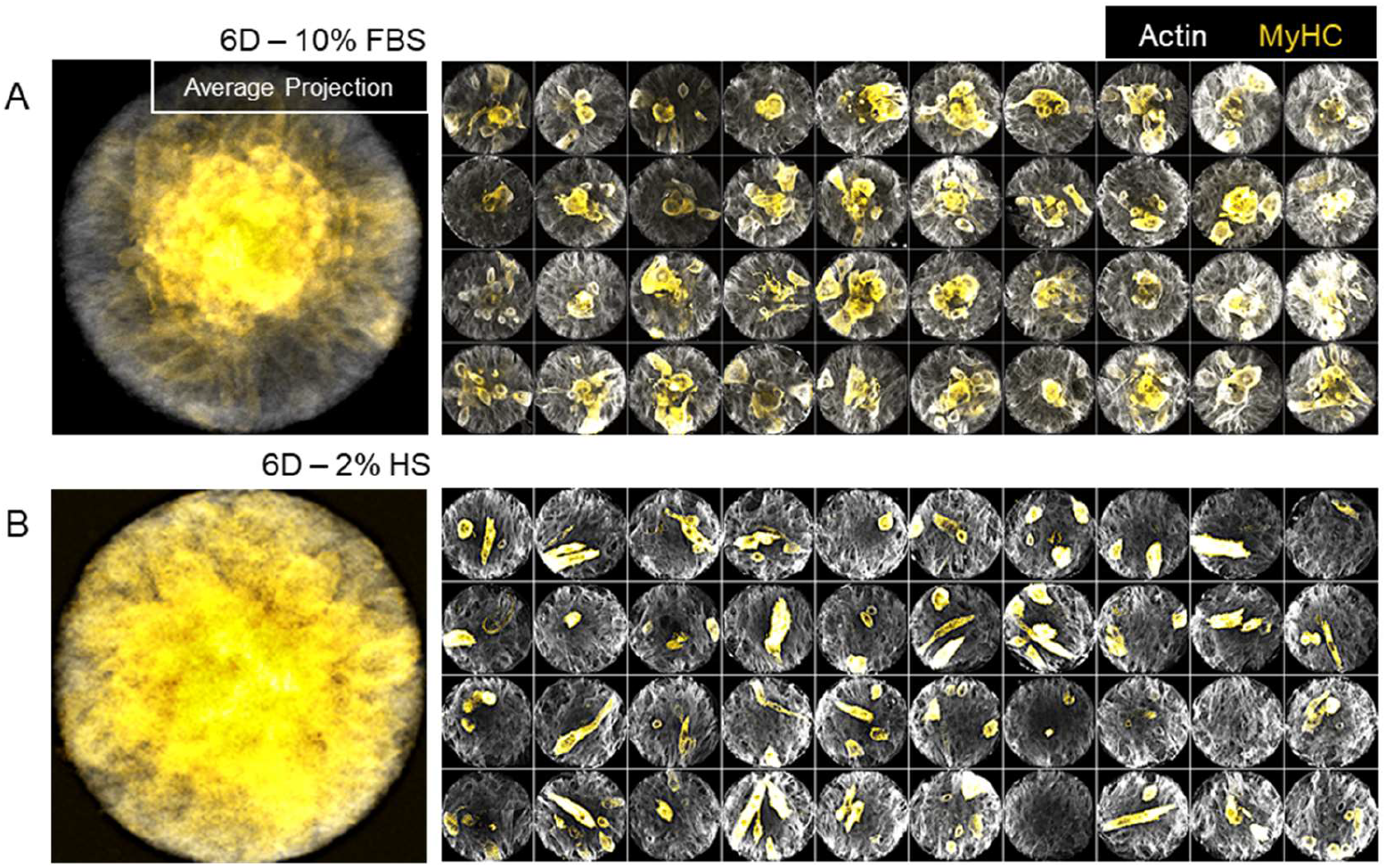
Localization of myosin heavy chain expression. (**A**) Average projection of confocal micrographs of cellular islands (N=40, r=100μm) grown for 6 days in complete medium (10% FBS). Individual micrographs are shown in the panels at the right. (**B**) Same analysis corresponding to cellular islands (N=40, r=100 μm) grown for 6 days under starvation conditions (2% HS).

**Figure S13.**
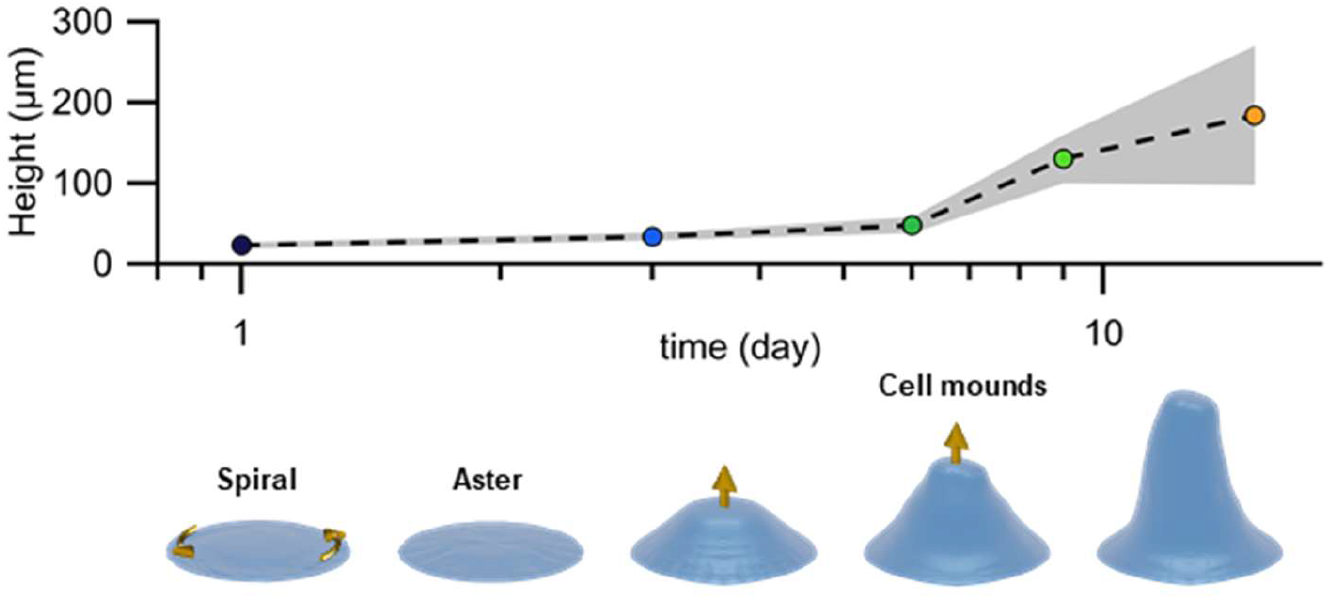
Growth of cellular mounds and protrusions. Average height of C2C12 islands at different timepoints after confluence (r=100μm).

## Supplementary movies

**Movie S1. Unconfined monolayer of C2C12 myoblasts.** Phase contrast time-lapse of a proliferating monolayer of myoblasts.

**Movie S2. Spiral-to-aster transition in a C2C12 myoblast disk.** Phase contrast time-lapse of myoblast monolayers under circular confinement. In time, cells rearrange from spiral arrangements into aster arrangements.

**Movie S3. Formation of cellular mounds.** Phase contrast time-lapse showing the formation of cellular mounds in the center of an aster of myoblasts.

**Movie S4. Cellular spirals.** Phase contrast time-lapse of low-density circular islands of myoblasts featuring spiral configurations. Division was blocked with Mitomycin-C.

**Movie S5. Actin dynamics in cellular spirals.** Fluorescence confocal time-lapse of the bottom plane of a cellular spiral. Actin was stained with SiR-Actin.

**Movie S6. Cellular asters.** Phase contrast time-lapse of high-density circular islands of myoblasts. Division was not blocked.

**Movie S7. Actin dynamics in cellular asters.** Fluorescence confocal time-lapse of the bottom plane of a cellular aster. Actin was stained with SiR-Actin.

**Movie S8. Pillar constriction experiment.** Differential interference contrast (DIC) time-lapse showing myoblasts constricting soft hydrogel pillars of different sizes. The consequent pillars’ deformation can be observed in the 3D renderings of the pillars obtained from fluorescence images’ segmentation.

**Movie S9. Dynamics and collapse of 3D cellular protrusions.** Phase contrast time-lapse of a myoblast protrusion, which collapses after confining pattern is degraded.

